# Peritoneal M2 macrophage-derived extracellular vesicles as natural multi-target nanotherapeutics to attenuate cytokine storm after severe infections

**DOI:** 10.1101/2022.03.13.484180

**Authors:** Yizhuo Wang, Shuyun Liu, Lan Li, Ling Li, Xueli Zhou, Meihua Wan, Peng Lou, Meng Zhao, Ke Lv, Yujia Yuan, Younan Chen, Yanrong Lu, Jingqiu Cheng, Jingping Liu

## Abstract

Cytokine storm is a primary cause for multiple organ damage and death after severe infections, such as SARS-CoV-2. However, current single cytokine-targeted strategies display limited therapeutic efficacy. Here, we report that peritoneal M2 macrophages-derived extracellular vesicles (M2-EVs) are multi-target nanotherapeutics to resolve cytokine storm. In detail, primary peritoneal M2 macrophages exhibited superior anti-inflammatory potential than immobilized cell lines. Systemically administrated M2-EVs entered major organs and were taken up by phagocytes (*e*.*g*., macrophages). M2-EVs treatment effectively reduced excessive cytokine (*e*.*g*., TNF-α and IL-6) release *in vitro* and *in vivo*, thereby attenuated oxidative stress and multiple organ (lung, liver, spleen and kidney) damage in endotoxin-induced cytokine storm. Moreover, M2-EVs simultaneously inhibited multiple key proinflammatory pathways (*e*.*g*., NF-κB, JAK-STAT and p38 MAPK) by regulating complex miRNA-gene and gene-gene networks, and this effect was collectively mediated by many functional cargos (miRNAs and proteins) in EVs. In addition to the direct anti-inflammatory role, human peritoneal M2-EVs expressed angiotensin-converting enzyme 2 (ACE2), a receptor of SARS-CoV-2 spike protein, and thus could serve as nanodecoys to prevent SARS-CoV-2 pseudovirus infection *in vitro*. As cell-derived nanomaterials, the therapeutic index of M2-EVs can be further improved by genetic/chemical modification or loading with specific drugs. This study highlights that peritoneal M2-EVs are promising multifunctional nanotherapeutics to attenuate infectious diseases-related cytokine storm.

## Introduction

Infectious diseases, caused by various types of pathogens (*e*.*g*., bacteria, viruses, fungi and parasites), have been a huge public health problem worldwide, which can affect ∼10% of all patients per year according to the World Health Organization report [1]. Although mild infections may be eliminated by the host immune system, severe infections frequently trigger immune disorders and hyper-inflammatory status, which is also known as cytokine storm [2, 3]. Cytokine storm, featured by excessive release of proinflammatory cytokines (*e*.*g*., IL-6 and TNF-α) and chemokines (*e*.*g*., MCP-1 and CCL5), has been recognized as one of the leading causes for development of multiple organ damage and failure after severe infections [4, 5]. For example, the recent COVID-19 pandemic caused by severe acute respiratory syndrome coronavirus 2 (SARS-CoV-2) is highly accompanied with elevated levels of cytokines (*e*.*g*., TNF-α and IL-6, ∼2-5 fold *vs*. normal healthy), which are mainly secreted by disordered immune cells (*e*.*g*., macrophages) in patients, and this effect is primarily responsible for the following acute respiratory distress syndrome and high mortality (up to ∼9.6%) [6, 7]. Therefore, therapeutics that can effectively resolve cytokine storm are urgently required in clinic. Some therapies aimed to counter or neutralize a single cytokine (*e*.*g*., IL-6 or TNF-α blockers) had been developed and shown certain beneficial effects in pre-clinical models[8, 9], but they showed controversial results in clinical trials of COVID-19 treatment[10]. The possible reason is that single cytokine-targeted therapy may well-function when this cytokine is the driving factor in this disease, while the mechanisms of cytokine storm are complicated and many cytokines are involved[11]. Instead, novel therapies that target multiple proinflammatory pathways might be more efficient, since the basis consensus is that cytokine storm is a result of disrupted networks involved multiple immune cells and cytokines.

Macrophages are the primary innate immune cells participating in host defense after pathogen infections, which distribute widely *in vivo* to sustain tissue homeostasis. In fact, the abnormal activation of macrophages has been linked to cytokine storm caused by severe infections and autoimmune diseases, which is also known as macrophage activation syndrome[12, 13]. Macrophages are heterogeneous immune cells and can be roughly divided into two subtypes: M1-like macrophages (M1-Mφ) and M2-like macrophages (M2-Mφ). M1-Mφ are mainly involved in the proinflammatory processes, and the hyper-activated M1-Mφ are the major source of numerous cytokines (such as IL-6 and TNF-α) to trigger tissue damage after infections[14]. In contrast, M2-Mφ mainly play anti-inflammatory and tissue repair roles. For example, we and others have found that M2-Mφ transplantation could reduce proinflammatory cytokine (*e*.*g*., IL-6 and TNF-α) production in mouse models of acute kidney injury and colitis[15]. However, as living cells, the transplantation of whole M2-Mφ *in vivo* is largely limited by several potential problems. For example, the viability and/or bioactivity of M2-Mφ may change by the storage and shipment conditions, and the phenotype of transplanted M2-Mφ may return to M1 due to the oxidative stress and inflammatory microenvironments *in vivo*. Thus, more stable, convenient and safe therapies are needed.

Extracellular vesicles (EVs) are a group of nanovesicles (∼30-1000 nm) secreted by living cells. EVs possess similar bioactive properties of donor cells due to the various types of biomolecules (proteins, nucleic acids, lipids, metabolites, *etc*.) they carried during biogenesis[14, 16]. For example, EVs from bone marrow-derived M2 macrophages reduced proinflammatory cytokines (*e*.*g*., IL-1β and IL-6) production in models of colitis and sepsis [17, 18]. As a type of natural nanovesicles released by cells, EVs exhibit several advantages, such as lower immunogenicity/toxicity and higher tissue penetrability when compared to whole cells and synthetic nanoparticles[16]. Moreover, EVs remain to be bioactive after long-term storage, repeated freeze/thaw cycles and other harsh handling conditions[16]. However, several questions need to be addressed before further translation of M2-EVs based therapies into clinical applications, especially in preventing infection-related cytokine storm. For example, the selection of proper EV resources (cell lines or primary cells), the therapeutic capacity of EVs in infectious models, and the remaining unclear comprehensive mechanisms of EVs.

Here, we report that primary peritoneal M2-EVs are efficient to reduce cytokine storm and its-related oxidative stress and multiple organs damage in mice challenged with bacterial endotoxin. M2-EVs carried many functional miRNAs/proteins and thus simultaneously inhibited multiple essential proinflammatory pathways by regulating complex miRNA-gene and gene-gene networks. In addition, human peritoneal M2-EVs expressed angiotensin converting enzyme 2 (ACE2) and could be served as nanodecoys to prevent SARS-CoV-2 viral infection *in vitro*, while this effect was absent in MSC-EVs and dexamethasone (Dex) (a corticosteroid anti-inflammatory drug used in clinic). Altogether, our results suggest that M2-EVs are promising multi-target nanotherapeutics to attenuate cytokine storm caused by severe infections, such as COVID-19.

## Materials and methods

### Cell culture

Primary mouse peritoneal macrophages (mMφ) were isolated from peritoneal dialysis model of 8-week-old male C57BL/6 mice as previously described[19]. The isolated macrophages were cultured in RPMI 1640 medium (Gibco, CA, USA) supplemented with 10% heat-inactivated fetal bovine serum (FBS, Gibco), penicillin (50 U/ml), streptomycin (50 μg/ml) and recombinant mouse macrophage colony-stimulating factor (mM-CSF, 20 ng/ml, Novoprotein). Primary mMφ and mouse macrophage cell lines (RAW264.7) were induced to M2 by incubating with recombinant mouse IL-4 and IL-13 (mIL-4/mIL-13, 20 ng/ml each, Novoprotein) proteins for 48 h. Human peritoneal macrophages (hPMφ) were isolated from dialysate of PD patients as previously described[20] and cultured in RPMI 1640 medium (Gibco) with 10% FBS (Gibco), penicillin (50 U/ml), streptomycin (50 μg/ml) and recombinant human M-CSF (hM-CSF, 20 ng/ml, Novoprotein). hPMφ were induced to M2 by incubating with recombinant human IL-4 and IL-13 (hIL-4/hIL-13, 20 ng/ml each, Novoprotein) proteins for 48 h. Patient dialysate sample collection and human cell experiments were conducted according to the National Institutes of Health (NIH) guidelines and approved by the Ethics Committee Biomedical Research, West China Hospital of Sichuan University (Permit NO. 20201019).

Mouse MSCs (mMSCs) were isolated from bone marrow of 2-week-old male C57BL/6 mice as described before[21], and human MSCs (hMSCs) were obtained from Sichuan Neo-life Stem Cell Biotech&Sichuan Stem Cell Bank (Chengdu, China). MSCs were cultured in the Dulbecco’s Modified Eagle Medium (DMEM, Gibco) in a humidified atmosphere at 37°C with 5% CO2 supplemented with 10% FBS (Gibco), penicillin (50 U/ml) and streptomycin (50 μg/ml), and cells with 3-4 passages were used for EV isolation.

### EV isolation and characterization

EVs-depleted medium was prepared by ultracentrifugation (120,000× g, 16 h, 4 °C) as described before[21]. For collection of conditioned medium, M2 or MSCs were washed by PBS and cultured in EVs-depleted medium for 48 h. EVs were isolated from culture medium of cells using a differential ultracentrifugation method. Briefly, medium was centrifuged at 300 × g for 15 min, 2,000 × g for 15 min at 4°C, followed by ultracentrifugation at 110,000 × g for 90 min at 4°C on an Optima XPN -100 ultracentrifuge with SW32Ti rotor (Beckman Coulter, Brea, CA, USA). The isolated EVs was washed with PBS and followed with a secondary ultracentrifugation (110,000 × g for 90 min at 4°C). The pure EVs pellets were resuspended in PBS and stored at - 80°C for further use.

The morphology of EVs was observed on a transmission electron microscopy (TEM, H-600, Hitachi, Ltd., Tokyo, Japan) as previously described[21]. The size distribution of EVs was analyzed using a Nanoparticle Tracking Analyzer (NTA, Particle Metrix, Meerbusch, Germany) as described before[21]. Western blotting was performed to detect the positive markers (anti-TSG101, anti-Alix, anti-HSP70) and negative marker (GM130) of EVs.

### Cellular uptake assay of EVs

M2-Mφ or mMSCs-derived EVs were labeled with PKH26 (Sigma Aldrich, St. Louis, MO, USA) according to the manufacturer’s protocols. mMφ cells were incubated with PKH26 labeled EVs (10 μg/ml) for 4 h at 37°C with 5% CO2. After incubation, cells were washed twice with PBS and stained with phalloidin (Yeasen Biotechnology, Shanghai, China) according to the manufacturers’ instructions. The stained cells were observed using a confocal laser scanning microscope (N-STORM &A1, Nikon, Tokyo, Japan). To determine the cellular uptake mechanism of M2-EVs, cells were pretreated with inhibitors including Bafilomycin A1 (10 nM, Glpbio Technology, Montclair, CA, USA), Cytochalasin D (0.5 μM, Glpbio Technology), and Wortmannin (0.5 μM, Glpbio Technology) for 30 min, and then incubated with PKH26-labeled EVs. After incubation for 4 h, cells were stained with phalloidin and observed using a confocal laser scanning microscope (Nikon).

### *In vitro* cytokine release model and EV treatments

Primary mMφ were treated with LPS (40 ng/ml, Novoprotein) and IFN-γ (20 ng/ml, Novoprotein) for 4 h at 37°C with 5% CO_2_ to induce cytokines release. For EV functional evaluation, M2-EVs or MSC-EVs (10, 20 μg/ml) were added at the same time with LPS/γ-IFN treatment. After incubation, the treated cells were collected for qPCR assay.

### *In vivo* biodistribution of EVs in mice

M2-Mφ or MSC-EVs were labeled with sulfo-Cy7-NHS ester according to the manufacturer’s instructions (New Research Biosciences Co., Ltd, Xi’an, China). The unbinding dyes were removed using an Exosome Spin Column (MW3000, Invirogen). Cy7-labeled M2-EVs or MSC-EVs (Cy7-EVs, 80 μg in 100 μl PBS per mouse) and an equal amount of free Cy7 (dye alone, negative control) was intravenously injected (*i*.*v*.) into C57/BL6 mouse *via* tail vein. At indicated time points after injection, mice were scarified by overdose of pentobarbital sodium anesthesia, and the organs including hearts, lungs, livers, kidneys, and spleens were collected and observed using an IVIS Spectrum optical imaging system (Perkin Elmer, Waltham, MA, USA). The intensity of fluorescence was quantified by Analyze 12.0 software (Perkin Elmer). To detect the distribution of EVs in organs, frozen sections of lung, liver, and spleen were made and co-stained with anti-CD68 (Abcam, 201340) and DAPI (Sigma). The digital images of stained sections were captured using a confocal laser scanning microscope (Nikon).

### *In vivo* model of cytokine storm and EV treatments

All animal experiments were conducted according to the National Institutes of Health (NIH) guidelines and approved by the Animal Care and Use Committee of West China Hospital, Sichuan University (Permit NO. 2021020A). Male C57BL/6 mice (∼20-25 g) were purchased from Experimental Animal Center of Sichuan University (Chengdu, China). Animals were housed in individual cages with controlled temperature, humidity, 12 h cycles of light and dark, and fed with standard chow and tap water *ad libitum*. Mice were randomly divided into four groups (n = 6∼9): healthy control (CON), LPS, LPS + M2-EVs, LPS + MSC-EVs. Mouse model of cytokine storm was induced by intraperitoneal (*i*.*p*.) injection with LPS (5 or 10 mg/kg, in 100 μl PBS, Sigma). For EV treatments, mice pre-treated with LPS were intravenously injected with M2-EVs or MSC-EVs (80 μg in 100 μl PBS per mouse) at 15 min after modeling. For detection of cytokine levels, serum and tissues were respectively collected from mice at 4 h after EV treatment. For analysis of immune cell populations, spleens were collected at 4 h after EV treatment. To evaluate the degree of organ damage, tissues including lung, liver, spleen and kidney were collected at 24 h after EV treatment.

### ELISA analysis of cytokine levels

The cytokines (TNF-α and IL-6) levels in cell culture medium or serum samples were analyzed using commercial ELISA kits for mouse TNF-α (Dakewei, Beijing, China) and mouse IL-6 (Dakewei) according to the manufacturer’s instructions. In brief, 50 µl of sample or standard was added to each well of a 96 -well plate, and then incubated with anti-TNF-α or anti-IL-6 at 37 °C for 90 min. Each well was incubated with streptavidin-HRP at 37 °C for 30 min. After incubating with TMB and stop solution, the 96-well plate was measured at 450 nm using a microplate reader (BioTek Instruments, Ink, USA).

### Western blot analysis

Cell samples or EV samples were lysed in radioimmunoprecipitation assay (RIPA) buffer supplemented with protease inhibitors (Calbiochem, CA, USA) and phosphatase inhibitors (Calbiochem). The total protein concentration was determined using a BCA Protein assay kit (CWBIO, Beijing, China). Equal amounts of proteins were electrophoresed on 12% sodium dodecyl sulfate-polyacrylamide gels (SDS-PAGE) and then transferred to polyvinylidene difluoride membranes (PVDF, Millipore, USA). PVDF membrane was blocked with 5% skim milk buffer and then incubated with one of the following primary antibodies: mouse anti-HSP70 (ab2787, Abcam, Cambridge, MA, USA), rabbit anti-TSG101 (14497-1-AP, Proteintech, Rosemont, IL, USA), rabbit anti-Alix (12422-1-AP, Proteintech), rabbit anti-GM130 (12480, Cell Signaling Technology, Beverly, MA, USA), rabbit anti-HGF (26881-1-AP, Proteintech), rabbit anti-IL-10 (20850-1-AP, Proteintech), rabbit anti-SOD2 (24127-1-AP, Proteintech), rabbit anti-TGF-β (21891-1-AP, Proteintech), mouse anti-ACE2 (66699-1-Ig, Proteintech), rabbit anti-Flag (AE004, ABclonal Biotech Co., Ltd., Cambridge, MA, USA) overnight at 4 °C. The PVDF membranes were incubated with HRP-conjugated secondary antibodies at 37 °C for 1 h. The protein bands on the PVDF membranes were observed using an enhanced chemiluminescence kit (Millipore) and quantified using Image J software (NIH, Bethesda, MD, USA).

### RNA extraction and real-time PCR assay

Total RNA was extracted using Trizol reagent (Gibco, Life Technologies) and reverse-transcribed into cDNA using a HiScreiptQ RT SuperMix kit (Vazyme Biotech, Nanjing, China). Real-time polymerase chain reaction (PCR) was performed on a CFX96 real-time PCR detection system (Bio-Rad, Hercules, CA, USA) with SYBR Green PCR mix (Vazyme Biotech). Primers used in this study are listed in Table S1. The data were analyzed using Bio-Rad CFX Manager software, and the relative changes of mRNA were calculated by delta-delta Ct method with GAPDH as the internal reference gene.

### Flow cytometric analysis (FCA)

The FCA samples of mouse spleen were prepared as previously reported[19]. In brief, mouse spleen tissues were ground in PBS and then passed through a 70 μm filter. Spleen samples were then incubated in red blood cell lysis buffer (R1010, Solarbio, Beijing, China) for 20 min to lyse red blood cell, followed by re-suspending the spleen sample with PBS. The prepared single-cell suspension from spleen was stained with FVS (564997, BD Pharmigen, San Diego, CA, USA), APC-Cy^™^7 Rat anti-mouse CD45 (557659, BD), PE Rat anti-mouse F4/80 (565410, BD), APC anti-mouse CD11c (117310, Biolegend, San Diego, CA, USA), FITC Hamster anti-mouse CD3e (553061, BD), PerCP/Cyanine5.5 anti-mouse CD4 (100434, Biolegend) and PE-Cy^™^7 Rat anti-mouse CD8a (552887, BD) for 30 min. After washing with PBS, the stained cells were analyzed using a flow cytometer (BD Bioscience, San Diego, CA, USA).

### Histological examination

Mouse tissues fixed in 4% paraformaldehyde were embedded in paraffin, and were sliced into 4 μm thick sections for hematoxylin and eosin (H&E) staining and immunohistochemistry (IHC) staining. For IHC, tissue sections were deparaffinized in xylene and rehydrated in graded ethanol, and the antigens were retrieved with citrate buffer. After inactivation of endogenous peroxidase with 3% H2O2, sections were blocked with 1% bovine serum albumin (BSA) and incubated with diluted rabbit anti-ICAM-1 (10831-1-AP, Proteintech) overnight at 4°C, and then stained with horse radish peroxide (HRP)-conjugated secondary antibodies (Abclonal) and 3,3-diaminobenzidine (DAB) substrate. Images of stained sections were captured with a light microscope (Zeiss, AX10 imager A2, Oberkochen, Germany) and quantified using the NIH Image J software.

### TUNEL staining of tissues

Terminal deoxynucleotidyl transferase dUTP nick end labeling (TUNEL) staining was carried out with commercial kit (Promega, Madison, WI, USA) following the manufacturer’s instructions, and nuclei were stained with DAPI (Sigma, USA). The stained lung, liver and spleen sections were observed under a fluorescence microscope. The average number of TUNEL^+^ cells in 10 fields of each section was quantified with fluorescence intensity using NIH Image J software.

### Immunofluorescent staining

After fixation (4% paraformaldehyde for 10 min), permeabilization (0.3% Triton X-100 for 10 min), and blocking (1% BSA for 1 h), cells or sections were incubated with mouse anti-8-OHdG (sc-393871, Santa Cruz Biotechnology, Dallas, TX, USA), anti-mouse KIM-1 (AF1817, R&D Systems, Minneapolis, MN, USA), anti-mouse CD68 (ab201340, Abcam) and anti-human ACE2 (266699-1-Ig, Proteintech) overnight at 4 °C. After washing with PBS, slides were incubated with fluorescein isothiocyanate (FITC)-conjugated secondary antibody (A11017, Life Technologies, CA, USA) at 37 °C for 1 h. Nuclei were visualized by staining with DAPI. Digital images were captured using a confocal laser scanning microscope (Nikon).

### mRNA sequencing analysis of cells

Total RNA of the cells was extracted using Trizol and then treated with DNase I (Takara, Shiga, Japan) to deplete the genomic DNA according the manufacturer’s instructions. High-quality RNA determination by 2100 Bioanalyser (Agilent, USA) and quantification by the ND-2000 (NanoDrop Technologies, USA) were used to construct sequencing library. RNA sequencing transcriptome library was prepared following TruSeq^™^ RNA sample preparation Kit from Illumina (San Diego, CA). cDNA was synthesized using a SuperScript double-stranded cDNA synthesis kit (Invitrogen) with random hexamer primers (Illumina). Libraries were size selected for cDNA target fragments of 300 bp on 2% Low Range Ultra Agarose followed by PCR amplification using Phusion DNA polymerase (New England Biolabs, Ipswich, MA, USA). The expression of each transcript was calculated according to the transcripts per million reads method. The PCA, volcano plot, and heatmap were generated using an online platform (https://www.omicsolution.org/wkomics/main/).

### miRNA sequencing analysis of M2-EVs

Total RNA of M2-EVs was extracted using Trizol and then treated with DNase I to deplete the genomic DNA. The Multiplex Small RNA Library Prep Set (New England Biolabs) was used to produce small RNA libraries. cDNAs were prepared using adaptor-specific primers, and DNA fragments (∼140-160 bp) were recovered. The library quality was evaluated using DNA High Sensitivity Chips on an Agilent Bioanalyzer 2100 system. Sequencing of libraries was conducted on an Illumina Novaseq 6000 system and 50 bp single-end reads were generated. All identical sequences with sizes of 18 to 32 nt were counted and removed from the initial data set. After elimination of non-miRNAs (rRNA, tRNA, snoRNA, *etc*.), the expression of miRNAs was analyzed using the perfectly matched sequences from the BLAST search of the miRbase (version 21.0). The cluster plot of miRNAs was generated by an online platform (https://www.omicsolution.org/wkomics/main/). The miRNA-gene interaction was analyzed using miRNet (https://www.mirnet.ca/)[22].

### miRNA inhibition assay

To determine the role of each miRNA in regulating cytokine production, PMφ was firstly transfected with individual miRNA inhibitors (10 nM, miR-21a-5p, miR-23b-5p, miR-342-5p, miR-378a-3p, miR-532-5p, miR-6238 or let-7b-5p) using Lipo6000 (C0526, Beyotime, Shanghai, China) for 4 h as previously reported[23]. After that, these cells were challenged with LPS/γ-IFN with or without M2-EVs (10 μg/ml) treatment. At indicated time after incubation, the changes in cytokines (TNF-α and IL-6) and proinflammatory pathways (NF-κB, JAK-STAT and p38 MAPK) of cells were analyzed.

### *In vitro* SARS-CoV-2 pseudovirus infection assay

Human embryonic kidney cells (HEK-293T cells) were transfected with human ACE2 lentivirus (MOI = 40, Obio Technology, Shanghai, China) in the presence of polybrene (5 μg/ml, Obio Technology), and the expression of ACE2 in HEK293 cells was detected by western bloting and IF staining. For SARS-CoV-2 pseudovirus infection assay, ACE2-overexpressed HEK-293T cells (2×10 ^4^ cells per well) were seeded in 48-well plate. SARS-CoV-2-GFP pseudovirus with lentiviral vector (MOI = 40, Obio Technology) were spinoculated with hM2-EVs (10 μg/ml), hMSC-EVs (10 μg/ml) or dexamethatsone (Dex, 0.5 μM) at 1200 g for 1.5 h at room temperature as previously reported[24], and the solution were then added into ACE2^+^ HEK-293T cells. At 48 h after infection, cells were washed, trypsinized and analyzed GFP expression using a flow cytometer (BD Bioscience).

### Statistical analysis

All data are presented as the mean ± SD and were analyzed with the t -test using GraphPad Prism 8.0.2 software (GraphPad Software Inc., San Diego, CA, USA); and p < 0.05 was considered statistically significant.

## Results and Discussion

### Primary M2 cells had stronger anti-inflammatory potency than immobilized cell lines

Macrophages are highly heterogeneous immune cells and widely distribute in different tissues *in vivo*, and their bioactivities may vary in different cell sources. For example, peritoneal macrophages infiltrated faster to the liver injury sites compared to bone marrow-derived macrophages, which then contributed to the local inflammation resolution[25]. According to previous studies, both primary macrophages (*e*.*g*., bone marrow-derived) and immobilized cell lines (*e*.*g*., mouse RAW264.7 cells) have been reported to produce M2-EVs. However, limited by the invasive surgery process and potential side effects, it is impossible to collect enough monocytes from the patients’ bone marrow by biopsy in clinical practice. Instead, it has been reported that large amounts of peritoneal monocytes/macrophages (∼40%) can be isolated from patients undergoing peritoneal dialysis (PD) [20]. Considering the superior yield and biofunctions, primary peritoneal Mφ (PMφ) were selected in this study. Although the anti-inflammatory effects of primary M2 and cell lines (*e*.*g*., RAW264.7)-derived M2 have been respectively reported[19, 26], whether they had comparable efficiency remains unclear. Therefore, we first isolated and polarized mouse peritoneal M2 cells and compared their anti-inflammatory capacity with RAW-M2 cells. As shown in Fig. S1A-B, the purity of isolated mouse PMφ was up to 95.7%. After induction with IL-4/IL-13, mouse PMφ and RAW264.7 cells could be polarized towards M2 subtype, as indicated by elevated gene expressions of Arg1, Mrc1 and CCL17 (Fig. S1C-D).

It is well-documented that the anti-inflammatory effects of M2 cells is largely mediated by paracrine mechanisms, such as the release of cytokines and EVs[14, 27]. Thus, the conditioned medium (containing cytokines and EVs) of cells was collected and used to evaluate their biofunction. As a key component of gram-negative bacterial outer membrane, lipopolysaccharides (LPS) is commonly used to mimic the bacteria-induced cytokine release *in vitro*[28]. LPS can be recognized by Toll-like receptors (TLRs) expressed on phagocytes (*e*.*g*., macrophages) and lead to the activation of downstream key proinflammatory pathways (*e*.*g*., MAPK and NF-κB) and the subsequent release of excessive cytokines[29]. In LPS/IFN-γ-primed macrophages, conditioned medium from M2-PMφ or M2-RAW cells decreased cytokine (TNF-α and IL-6) expressions in a dose-dependent manner (Fig. S2A). However, M2-PMφ group showed stronger inhibitory effect (∼2-fold, *vs*. M2-RAW264.7 group) on cytokine expressions in LPS/IFN-γ-primed macrophages (Fig. S2A). The production of key immunoregulatory factors, such as TGF-β[30], was also much higher in M2-PMφ than M2-RAW cells (Fig. S2B). These results suggest that the primary peritoneal M2 cells have stronger anti-inflammatory potency than immobilized cell lines, and they are more proper cell resources for M2-EVs production.

### M2-EVs effectively suppressed endotoxin-induced cytokine release *in vitro*

Next, we isolated EVs from culture medium of primary cells and analyzed their properties. The isolated M2-EVs and MSC-EVs had a typical bi-layer lipid membrane structure (Fig. 2A), and their average sizes were ∼150 nm and ∼170 nm, respectively (Fig. 2B). These EVs were positive for marker proteins (Alix, TSG101 and HSP70) and negative for GM130 (a Golgi marker) expression (Fig. 2C). EVs can regulate the function of target cells by delivering bioactive cargos, and thus the uptake of EVs by recipient cells is an essential step. Previous studies had found that the *i*.*v*. injected EVs were mainly taken up by phagocytes (*e*.*g*., macrophages) *in vivo*[31]. Therefore, we first assayed whether M2-EVs could enter into PMφ and explored the possible uptake mechanism *in vitro*. M2-EVs were labeled with PKH26 (a lipophilic fluorescent dye), and then incubated with primary PMφ for 4 h. After that, positive signals of EVs (red) were observed in the cytoplasm of phalloidin-labeled cells (green), suggesting that M2-EVs were taken up by PMφ (Fig. 2E). The mechanisms of EV uptake by recipient cells are complicated and multiple pathways involved, such as macropinocytosis, endocytosis, and phagocytosis, were all reported to play a part in EVs uptake[32, 33].

**Fig. 1.**
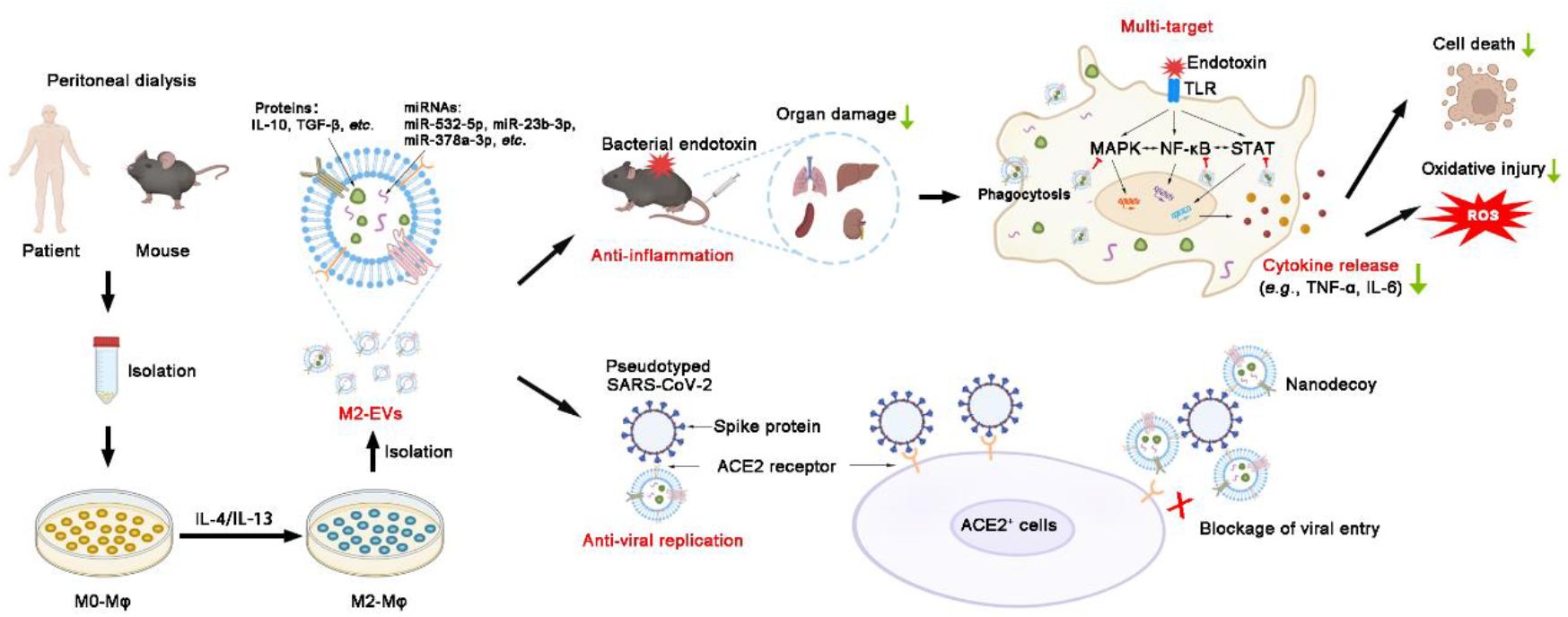
Schematic diagram of the findings of this study. Large amounts of macrophages can be isolated from peritoneal dialysates of patients and mice. M2-EVs from peritoneal M2 macrophages carry many functional cargos and thus efficiently attenuate endotoxin-induced cytokine storm and organ damage by inhibiting multiple essential proinflammatory pathways. Moreover, human M2-EVs can also serve as nanodecoys to block viral entrance and replication in cells due to expression of ACE2 (a receptor of SARS-CoV-2 spike protein).

**Fig. 2.**
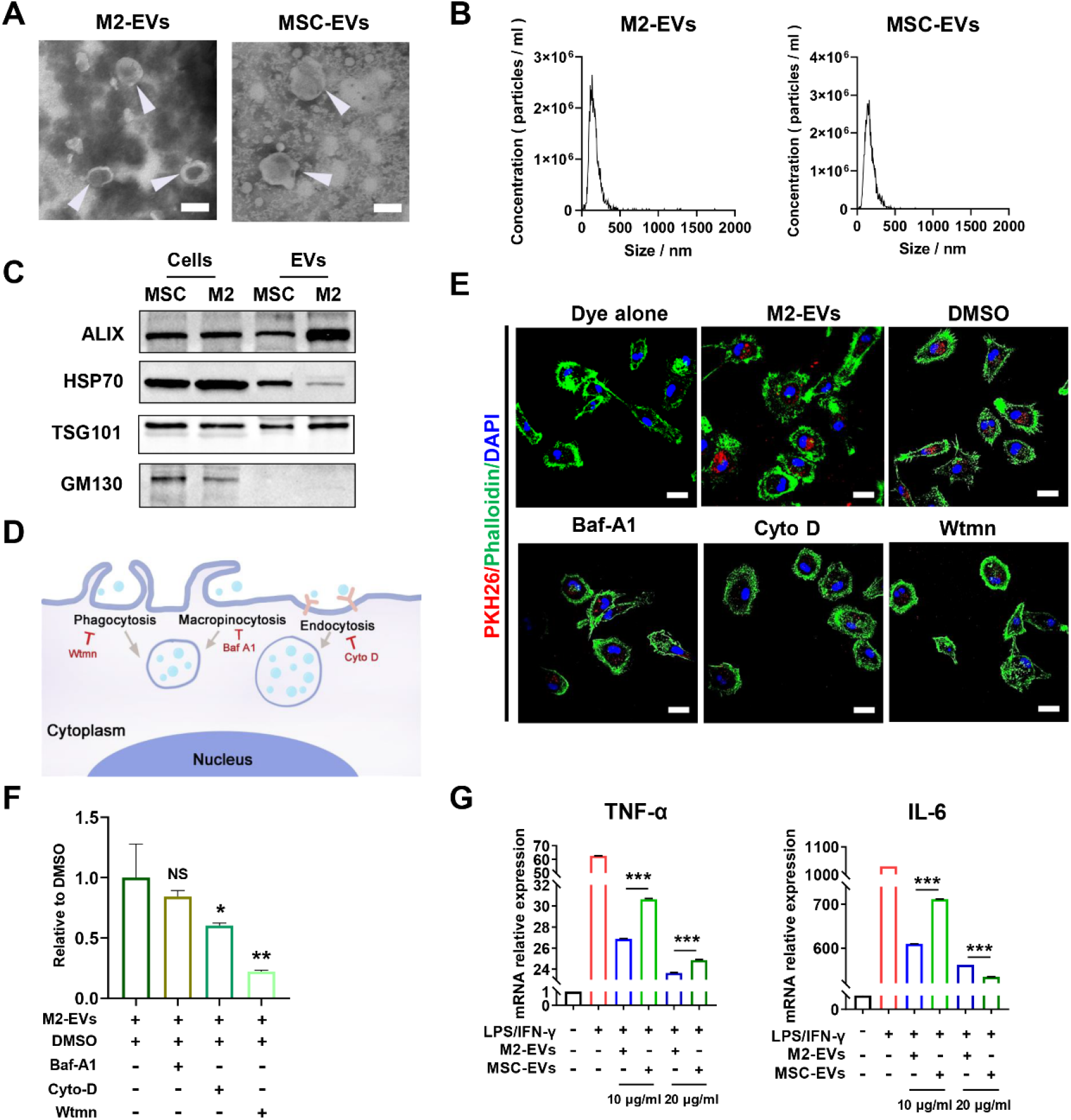
Isolation and evaluation of properties of M2-EVs *in vitro*. (A) Representative TEM images of EVs. White arrow indicated EVs (scale bar = 200 nm). (B) Size distributions of EVs measured by NTA. (C) Western blot analysis of EVs positive markers (ALIX, HSP70, TSG101) and negative markers (GM130). (D) Schematic illustration of the mechanisms of EVs uptake. (E) Representative images of M2-EVs uptake in PMφ stained with FITC Phalloidin (Green) and DAPI (Blue) (scale bar = 20 μm). Cells were pre-treated with DMSO, Baf-A1 (Bafilomycin A1), Cyto D (Cytochalasin D) or Wtmn (Wortmannin) for 15 min, and the incubated PKH26-labeled M2-EVs (Red) for 4 h. (F) Relative uptake ratio of M2-EVs in PMφ of different groups (n = 3; ^*^p < 0.05, ^**^p < 0.01 *vs* DMSO group). (G) Measurement of TNF-α and IL-6 mRNA levels in LPS/IFN-γ-primed PMφ with or without EVs treatments for 4 h by qPCR (n = 3; ^***^p < 0.001, M2-EVs *vs* MSC-EVs).

To explore the possible pathways that mediate M2-EVs uptake, PMφ were pre-treated with specific inhibitors including bafilomycin A1 (Bal-A1, a macropinocytosis inhibitor), cytochalasin D (Cyto-D, an endocytosis inhibitor) and wortmannin (Wtmn, a phagocytosis inhibitor) as previously reported[32, 34] (Fig. 2D). Compared to vehicle (DMSO) group, the signals of M2-EVs (red) were decreased in Bal-A1, Cyto-D and Wtmn treatment groups, and the lowest signal of M2-EVs was observed in Wtmn treatment group (Fig. 2E-F). These results suggested that the uptake of M2-EVs in PMφ was regulated by multiple pathways, and phagocytosis might be one of main routes.

The inhibitory effects of the primary M2-EVs on cytokine release were further assayed *in vitro*, and another type of well-studied MSC-EVs was chosen as the positive control, since MSC-EVs therapies have been widely reported to reduce cytokine release in various inflammatory and infectious disease models[35, 36]. Among many cytokines, the dramatic elevation of TNF-α and IL-6 has been most frequently reported in patients or animal models with cytokine storm. For instance, recent studies indicated that elevated IL-6 is highly associated with the increased mortality in COVID-19 patients[10]. IL-6 is an activator of key proinflammatory pathways (*e*.*g*., JAK-STAT) and can induce the expression of many other cytokines (*e*.*g*., TNF-α, IL-8) as well as the differentiation of multiple immune cells (*e*.*g*., B cells and T cells)[37]. TNF-α is a vital acute phase protein induced by infections, which can activate proinflammatory NF-κB pathway and act as an amplifier in inflammatory cascade[3]. Increased TNF-α is positively correlated with mortality in disease conditions, such as lethal toxic shock[38]. In this study, we found that LPS/IFN-γ priming induced a dramatic increase in TNF-α (∼60-fold) and IL-6 (∼1000-fold) gene expression in macrophages compared to control group (Fig. 2G). Conversely, M2-EVs treatment markedly decreased the expressions of TNF-α (decreased by ∼70%, *vs*. LPS/IFN-γ group) and IL-6 (decreased by ∼60%, *vs*. LPS/IFN-γ group) in a dose-dependent manner. MSC-EVs group also showed considerable reductions in TNF-α and IL-6 expressions, but their inhibitory efficacy was lower than M2-EVs group at the same concentrations (Fig. 2G). These results suggest that M2-EVs can suppress endotoxin-induced excessive cytokine release *in vitro*, and their therapeutic effects may be superior than MSC-EVs.

### M2-EVs attenuated cytokine storm and macrophage hyper-activation *in vivo*

EVs can affect the functions of distal cells or tissues which are far from where they are secreted or administrated. To visualize the biodistribution of systemically infused EVs, EVs were labeled with cyanine7 (a near-infrared dye, Cy7-EVs) and were then injected into mice *via* tail vein. In line with previous reports, we found that M2-EVs (Fig. 3A, C) and MSC-EVs (Fig. 3B, D) displayed a similar distribution pattern, which primarily accumulate in liver, spleen, lung, kidney and heart after *i*.*v*. injection. It has been reported that *i*.*v*. injected EVs are mainly taken up by phagocytes (*e*.*g*., macrophages) *in vivo*[39]. Consistently, we observed the colocalization of Cy7-labeled M2-EVs with the CD68^+^ macrophages in tissue sections, such as lung, liver and spleen (Fig. 3E). However, no detectable signal was observed in dye control (Cy7 alone) groups (Fig. 3A-E). These results suggest that large amounts of M2-EVs can arrive at major organs and are taken up by macrophages after systemic injection.

**Fig. 3.**
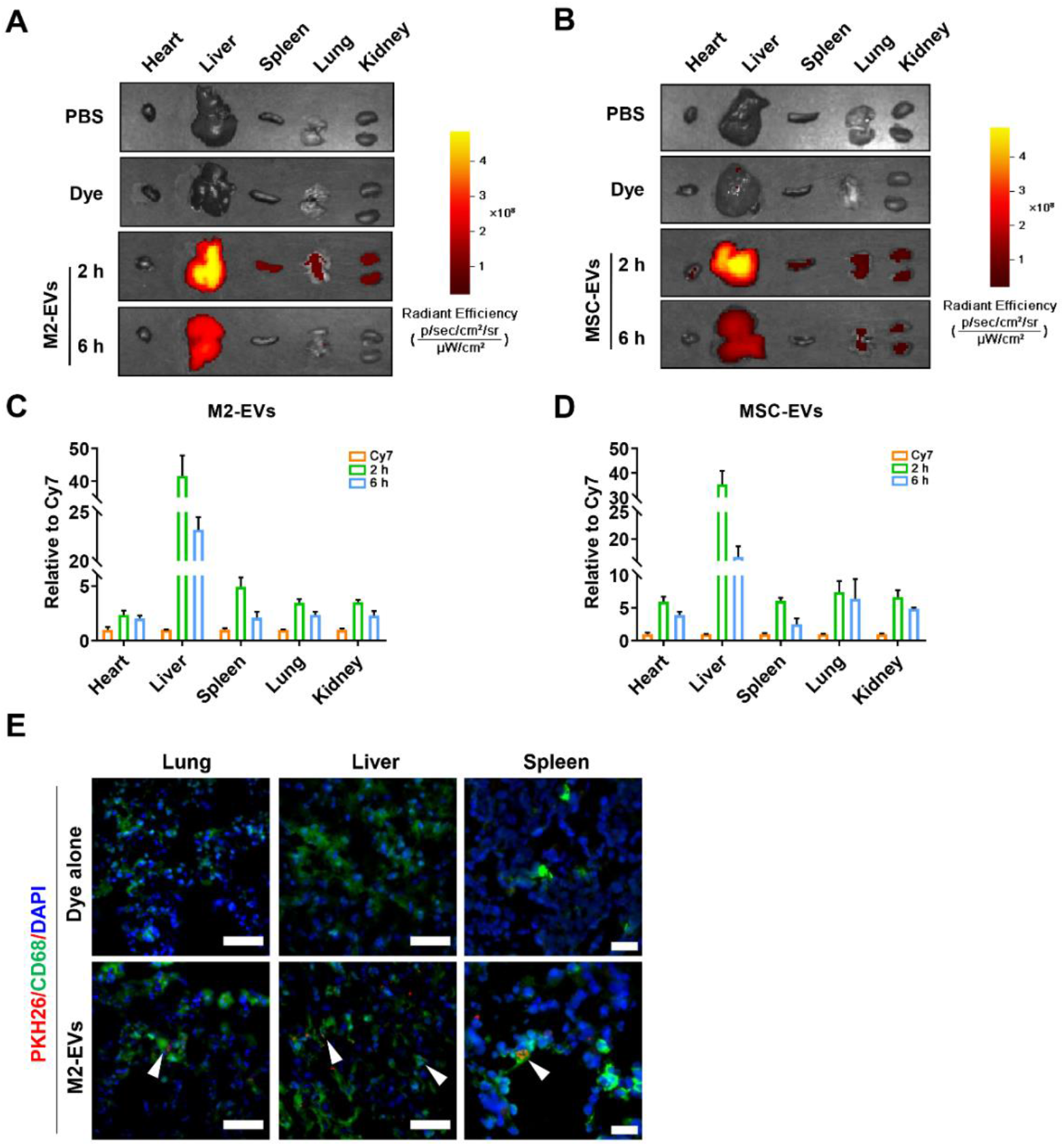
*In vivo* biodistribution of EVs. (A-B) Representative IVIS images of organs harvested from mice at 2 and 6 h after intravenous injection of Cy7-labeled M2-EVs or MSC-EVs. Mice received PBS and Cy7 dye alone were included as negative controls. (C-D) Quantification of fluorescent intensity of M2-EVs or MSC-EVs in different organs of mice (n = 3). (E) Representative micrographs of Cy7-EVs (Red) in tissue sections. Spleen tissues were co-stained with CD68^+^ macrophages (Green) and DAPI (Blue). White arrow indicated the co-localization of Green and Red signals (scale bar = 50 μm for lung and liver, scale bar = 20 μm for spleen).

To validate the *in vitro* findings, we also evaluated the therapeutic effects of EVs in a mouse model of LPS-induced cytokine storm (Fig. 4A). As a type of bacterial-derived endotoxin, LPS is a robust bacterial inducer of cytokine release *in vivo*, and low-dose injection of LPS into healthy human volunteers causes pathophysiologic alterations which are similar to those reported in patients with cytokine storm[40]. To better stimulate the severe infections, we first detected the dynamic changes in blood cytokine levels of mice in response to different doses of LPS (5 or 10 mg/kg) challenge. Higher dose of LPS (10 mg/kg) priming leads to a rapid and dramatic elevation of plasma TNF-α and IL-6 levels compared to control or lower dose (5 mg/kg) group (Fig. S3A-B). In line with the *in vitro* results, M2-EVs treatment effectively reduced the systemic and local inflammatory degree *in vivo*, as indicated by the reduction in plasma cytokines (decreased by ∼78% for TNF-α, decreased by ∼65% for IL-6) and tissue cytokines (decreased by ∼50-70% for TNF-α, decreased by ∼60-80% for IL-6) levels of multiple organs (liver, lung and kidney) compared to LPS groups (Fig. 4B-C). Moreover, the expression of vital chemokines (*e*.*g*., ICAM-1) was suppressed by M2-EVs treatment in lung, spleen and liver tissues (Fig. 4F, G). Similarly, we also observed a slightly higher anti-inflammatory efficacy in M2-EVs groups compared to MSC-EVs groups at same concentrations (Fig. 4B, C, F, G).

**Fig. 4.**
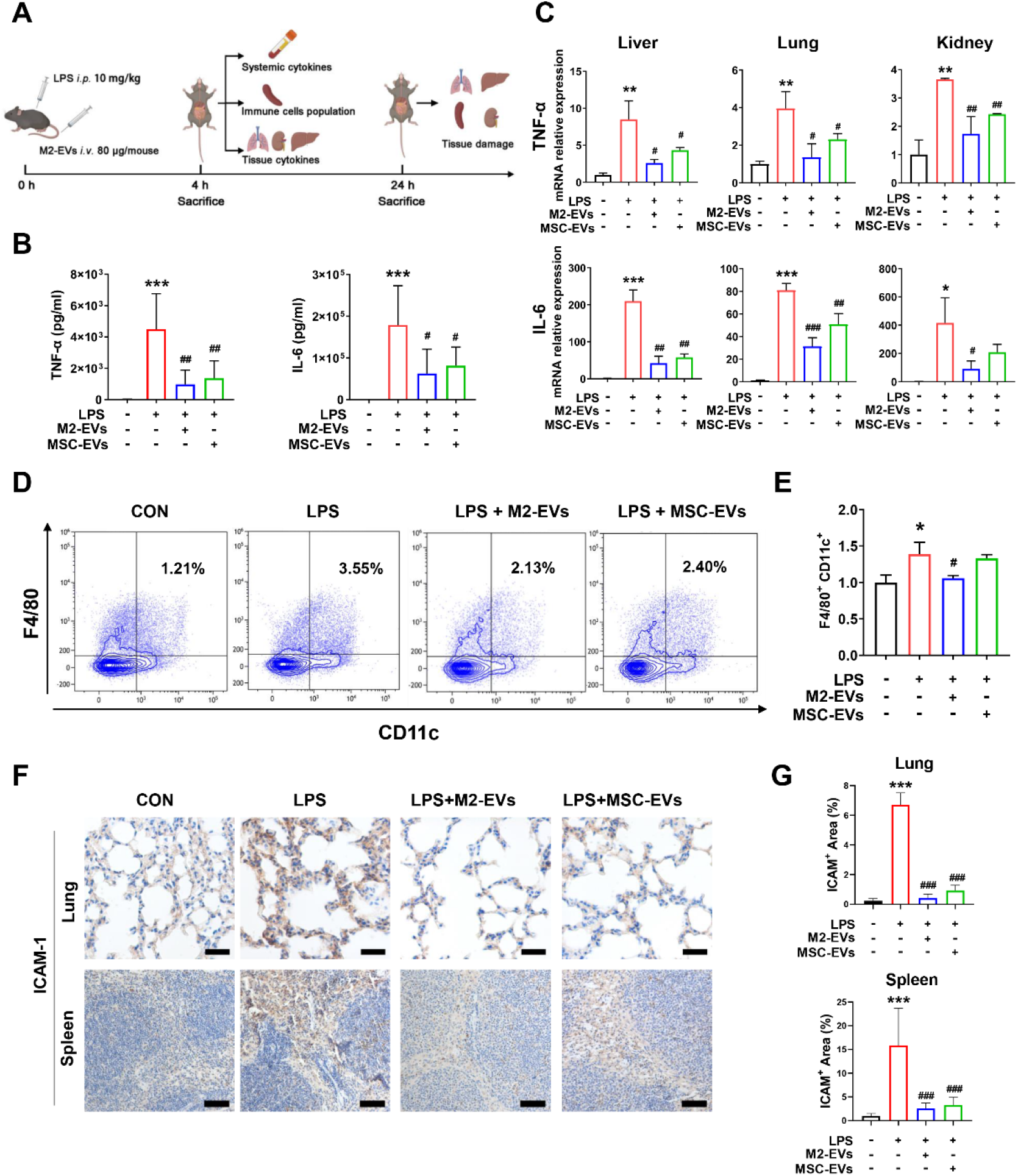
Effects of M2-EVs on inflammatory response *in vivo*. (A) Schematic illustration of animal experiments. (B) Measurement of mouse plasma TNF-α and IL-6 levels at 4 h after LPS challenge by ELISA (n = 6∼9). (C) Quantification of mouse TNF-α and IL-6 mRNA levels in liver, lung and kidney tissues at 4 h after LPS challenge by qPCR (n = 3). (D) FCA analysis of immune cells population changes in spleen. (E) Quantification of pro-inflammatory F4/80^+^/CD11c^+^ macrophages population in spleen (n = 3). (F) Representative micrographs of ICAM-1 IHC staining of lung and spleen sections at 24 h after LPS challenge. (G) Quantification of tissue ICAM-1 protein expression detected by IHC staining (n = 3). (^***^p<0.001, ^**^p < 0.01, ^*^p<0.05 *vs* CON group; ^###^p<0.001, ^##^p<0.01, ^#^p<0.05 *vs* LPS group; scale bar = 20 μm for lung; Scale bar = 100 μm for spleen).

In infectious conditions, the overproduction of cytokines are mainly regulated by a complex network involving innate and adaptive immune cells, such as monocytes, macrophages, T cells and dendritic cells[41], and the populations of each cell type dynamically change in different stages or states after infections. To further explore the immunoregulatory role of M2-EVs, we detected the changes in the immune cell populations in organs (*e*.*g*., spleen) using a multiple staining flow cytometry analysis (FCA) method (Fig. S4, Fig. 2D). Indeed, the population of proinflammatory M1 macrophages (F4/80^+^/CD11c^+^) increased markedly in spleen at the early stage (4 h) after LPS challenge, while its population was reduced by M2-EVs treatment (Fig. 4D-E). MSC-EVs treatment led to a slight reduction in proinflammatory macrophages (F4/80^+^/CD11c^+^) in spleen (Fig. 4D, E). However, there was no significant difference between the populations of CD45^+^ total leucocytes, CD3^+^ total T cells, CD4^+^ helper T cells and CD8^+^ cytotoxic T cells in the spleen tissues of mice at 4 h after LPS priming (Fig. S4B), suggesting that innate immune cells (*e*.*g*., macrophages) might be a major resource of cytokines at early phase after endotoxin challenge. These results suggest that M2-EVs can effectively reduce the endotoxin-induced cytokine storm *in vivo*, and this effect is at least partly induced by inhibiting hyper-activation of macrophages.

### M2-EVs attenuated cytokine storm-associated multiple organ damage *in vivo*

Infection-induced cytokine storm is highly associated with multiple organ damage, such as acute lung, liver and kidney injury [42]. Thus, we also determined whether M2-EVs treatment could protect multiple organs against inflammatory injury *in vivo*. After LPS challenge, we found clear pathological lesions, such as hemorrhage and inflammatory cell infiltrations in multiple organs (lung, liver and spleen) (Fig. 5A). Lung edema and thickened alveolar walls, liver necrosis, and dilated spleen sinus with a disorganized white pulp with loss in boundary definitions and barely distinct follicular structure, were also observed in mice with LPS challenge (Fig. 5A). The levels of kidney injury molecule-1 (KIM-1, a marker of renal damage) was elevated in kidneys from LPS group (Fig. S5A-B). In contrast, M2-EVs treatment suppressed these lesion formations in multiple organs, and reduced the overall tissue injury scores of lung, liver and spleen in mice with LPS priming (Fig. 5B, S5B). Meanwhile, MSC-EVs also exhibited similar effects on tissue protection *in vivo*.

**Fig. 5.**
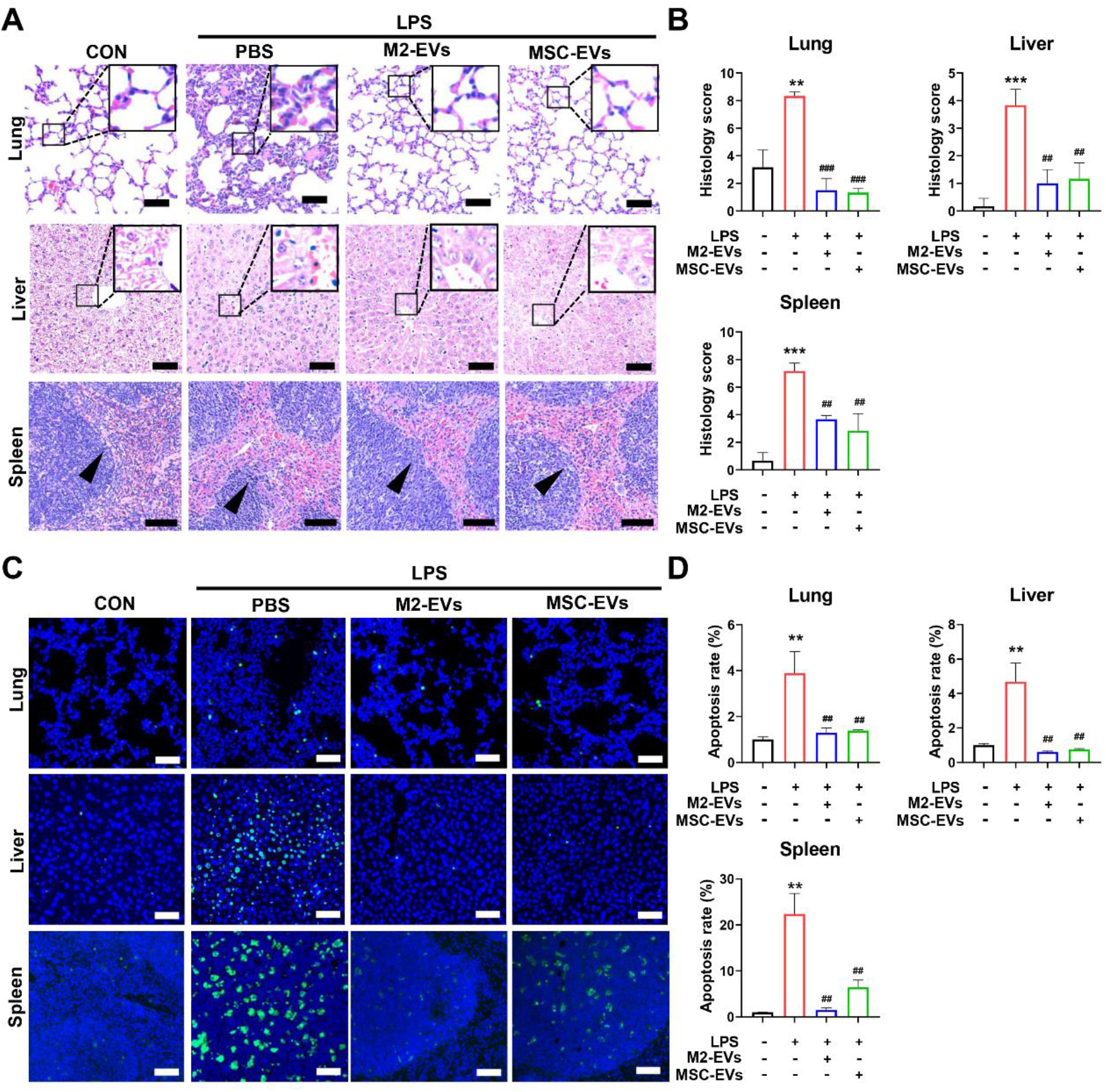
Effects of M2-EVs on multiple organ damage *in vivo*. (A) Representative H&E staining micrographs of the lung, liver and spleen sections at 24 h after LPS challenge (scale bar = 20 μm for lungs and livers; scale bar = 100 μm for spleens). The enlarged figures show the detailed changes in lung and liver sections and the black arrows in splenic micrographs showed the loss in boundary definitions. (B) Quantification of histology injury scores of lung, liver and spleen by professional pathologists (n = 3). (C) Representative TUNEL staining micrographs of the lung, liver and spleen sections at 24 h after LPS challenge (scale bar = 100 μm). (D) Quantification of TUNEL^+^ cells in the lung, liver and spleen sections. (n = 3). (^***^p< 0.001, ^**^p < 0.01 *vs* CON group; ^###^p<0.001, ^##^p<0.01 *vs* LPS group).

Excessive cytokines can induce cell death by triggering oxygen reactive species (ROS) overproduction and pro-apoptotic signals, which in turn cause severe tissue damage and final organ failure[43]. Moreover, cytokines and oxidative stress often contribute to fuel each another *via* complicated feedbacks [21, 44]. For example, TNF-α stimulation led to elevated mitochondrial ROS (mtROS) levels in the lung epithelial cells[45]. In line with these reports, we found an increase in mtROS production of PMφ under LPS/γ-IFN stimulation, and their levels were further reduced by M2-EVs treatment (Fig. S6A). The levels of 8-hydroxydeoxyguanosine (8-OHdG, an oxidative injury marker) increased in the major organs (lung, liver and spleen) of mice at 24 h after LPS challenge, which were neutralized by M2-EVs treatment (Fig. S6B). The apoptotic cells in tissues of mice were also detected by terminal deoxynucleotidyl transferase dUTP nick end labeling (TUNEL) staining. Consequently, we observed elevated numbers of TUNEL^+^ apoptotic cells in the major organs (lung, liver and spleen) of mice received LPS, whereas this elevation was markedly ameliorated by M2-EVs treatment (Fig. 5C-D).

### M2-EVs suppressed cytokine production by inhibiting multiple proinflammatory pathways

To further reveal the underlying mechanisms, we analyzed the changes in gene profiles of proinflammatory Mφ receiving M2-EVs treatment using RNA sequencing (RNA-seq). Principal component analysis (PCA) results showed an overall alteration in genes expression between groups (Fig. 6A). Compared with control group, 3008 upregulated genes and 2211 downregulated genes were observed in LPS/IFN-γ group. After M2-EVs treatment, 543 upregulated genes and 732 downregulated genes were found compared to LPS/IFN-γ group (Fig. 6B). Heatmap analysis showed that many upregulated genes induced by LPS/IFN-γ were neutralized by M2-EVs treatment (Fig. 6C). Next, we performed gene-gene interaction analysis and KEGG pathway enrichment analysis to reveal the intracellular pathways affected by M2-EVs. Indeed, these differentially expressed genes between the LPS/IFN-γ group and the LPS/IFN-γ + M2-EVs group were mainly enriched in inflammation-associated pathways, such as cytokine-cytokine receptor interaction, JAK-STAT, MAPK, and NF-κB pathways (Fig. 6D-E). Further western bloting results confirmed the activation of NF-κB, p38 MAPK and JAK-STAT pathways in LPS-treated macrophages, while these pathways were inhibited by M2-EVs treatment (Fig. 6F, 7D). These results suggest that, instead of targeting a single cytokine, M2-EVs may suppress cytokine production by simultaneously inhibiting multiple essential proinflammatory pathways (*e*.*g*., NF-κB, p38 MAPK and JAK-STAT).

**Fig. 6.**
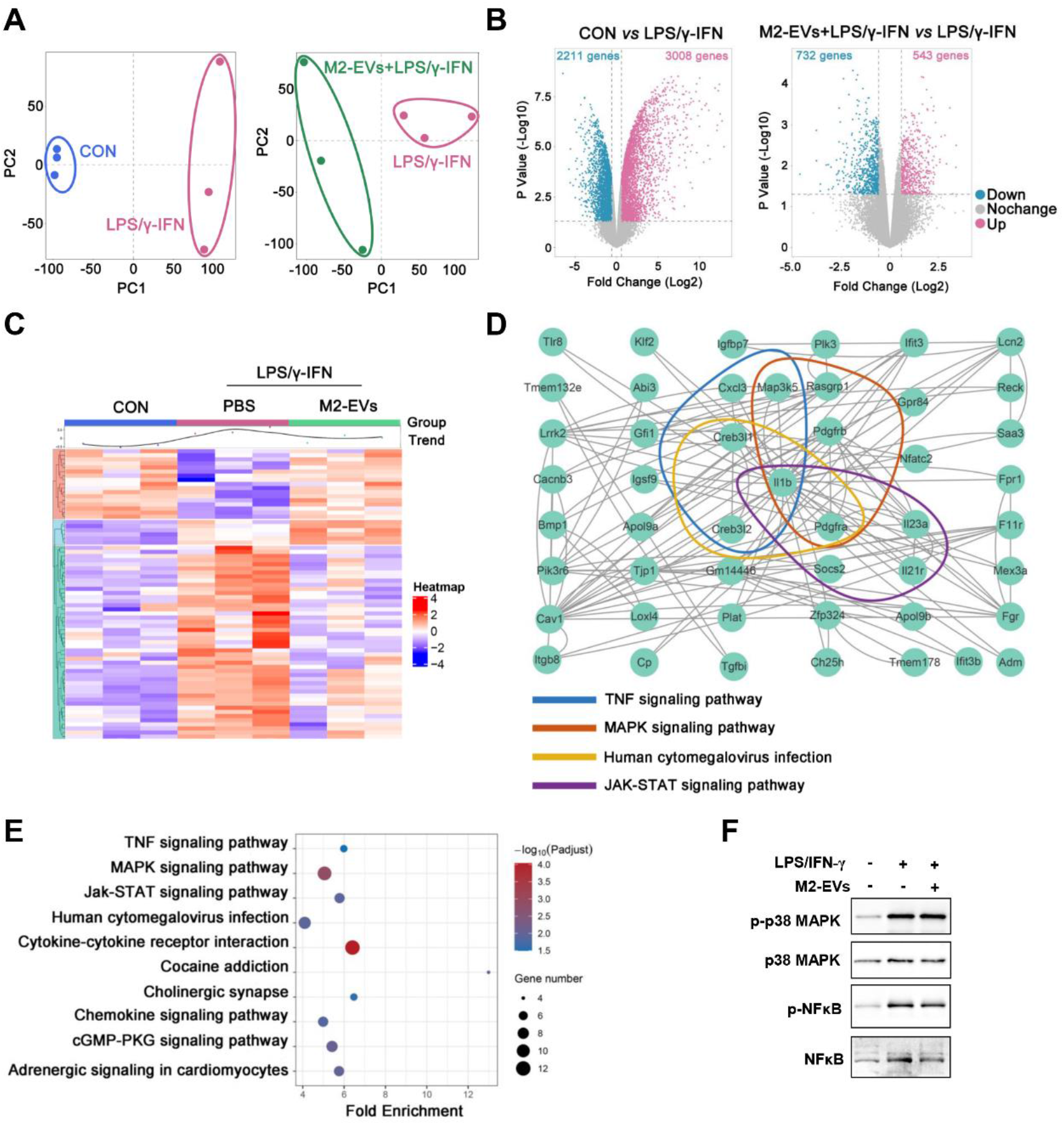
Effects of M2-EVs on proinflammatory pathways *in vitro*. (A) PCA score plot representing discrepancies between groups (n = 3). (B) Volcano plot representing the significantly changed genes (fold change > 2, p < 0.05). (C) Heatmap of the differentially expressed genes and the average expression curve above the heatmap. (D) Gene-gene interaction between the top 20 differentially expressed genes using String analysis. (E) KEGG pathway enrichment analysis based on the top 20 differentially expression of genes. (F) Western blot analysis of the total and phosphorylated levels of p38MAPK and NF-κB in LPS/IFN-γ-primed PMφ with M2-EVs treatment for 1 h.

Increasing evidence indicates that EVs can impact the functions of recipient cells by delivering multiple types of bioactive cargos, especially microRNAs (miRNAs) and proteins[14]. To explore the possible cargos responsible for the therapeutic roles of M2-EVs, we first detected the miRNA profile in M2-EVs by RNA-seq. A total of 601 miRNAs (including 445 known and 156 predicted miRNAs) were found in M2-EVs, and the top 8 miRNAs were having the most abundance, such as miR-342-3p, miR-674-5p, miR-24-3p, miR-21a-5p, miR-23b-3p, miR-146a-5p, miR-378a-5p and let-7b-5p were shown (Fig. 7A). Next, we performed miRNA-gene network analysis to explore the possible interactions between these miRNAs carried by EVs and the intracellular genes regulated by M2-EVs (Fig. 7B). Again, the results indicated that these miRNAs might jointly regulate gene networks mainly enriched in proinflammatory pathways, such as cytokine-cytokine receptor interaction, JAK-STAT, MAPK, NF-κB, TNF and TGF-β pathways (Fig. 7B). We further validated these findings using miRNAs inhibitors, which can specifically bind to endogenous miRNAs and thus inhibit their bioactivities. Indeed, the inhibitory effects of M2-EVs on cytokines (TNF-α and IL-6) production and proinflammatory pathways (NF-κB, JAK-STAT3 and p38 MAPK) were abolished to some extent after pre-treatment with individual miRNAs inhibitors, such as miR-532-5p, miR-23b-3p, let-7b-5p, miR-342-5p and miR378a-3p (Fig. 7C-E). In fact, these miRNAs (*e*.*g*., miR-532-5p, miR-23b-3p and miR-378a-3p), had been shown to reduce cytokine release in inflammatory disease models [46, 47]. Thus, our results suggest that the anti-inflammatory effects of M2-EVs were at least partly dependent on multiple functional miRNAs carried by these EVs.

**Fig. 7.**
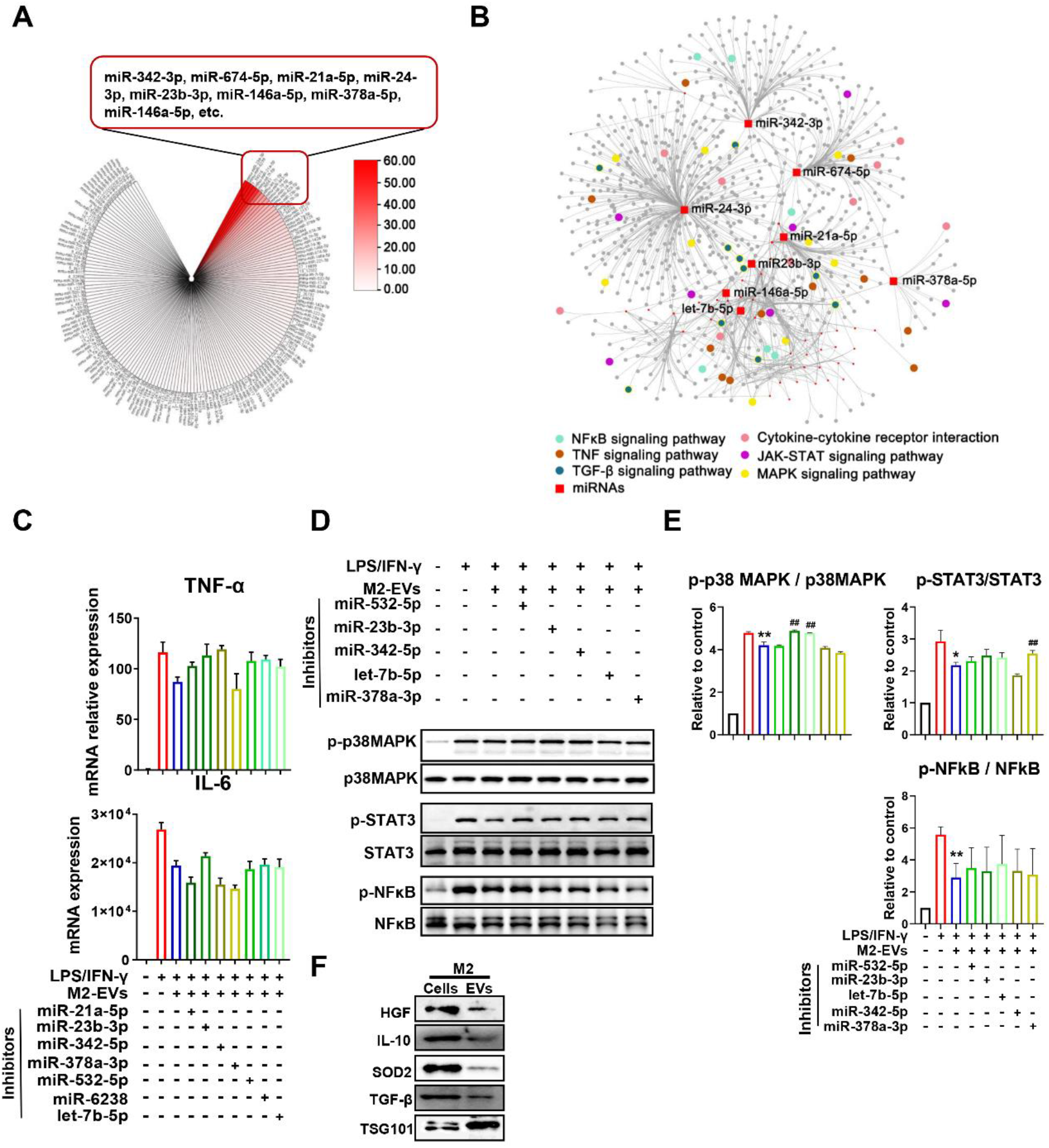
M2-EVs inhibited cytokine production by mediating complex signaling networks. (A) An overview of miRNAs contents within M2-EVs, and top 8 abundant miRNAs were listed. (B) miRNAs-genes network analysis and KEGG pathway enrichment analysis based on differently expressed genes. (C) Measurement of cytokine TNF-α and IL-6 mRNA levels in LPS/IFN-γ-primed PMφ co-treatment with M2-EVs and different miRNA inhibitors for 4 h. (D) Western blot analysis of the phosphorylation levels of p38MAPK, STAT3 and NF-κB in LPS/IFN-γ-primed PMφ co-treatment with M2-EVs and miRNA inhibitors for 1 h. (E) Quantification of the phosphorylation levels of p38MAPK, STAT3 and NF-κB in different group (n = 3; ^**^p < 0.01, ^*^p<0.05 *vs* CON group; ^##^p<0.01 *vs* LPS group). (F) Western blot analysis of HGF, IL-10, SOD2, TGFβ1 and TSG101 protein expressions in M2 cells and EVs.

**Fig. 7.**
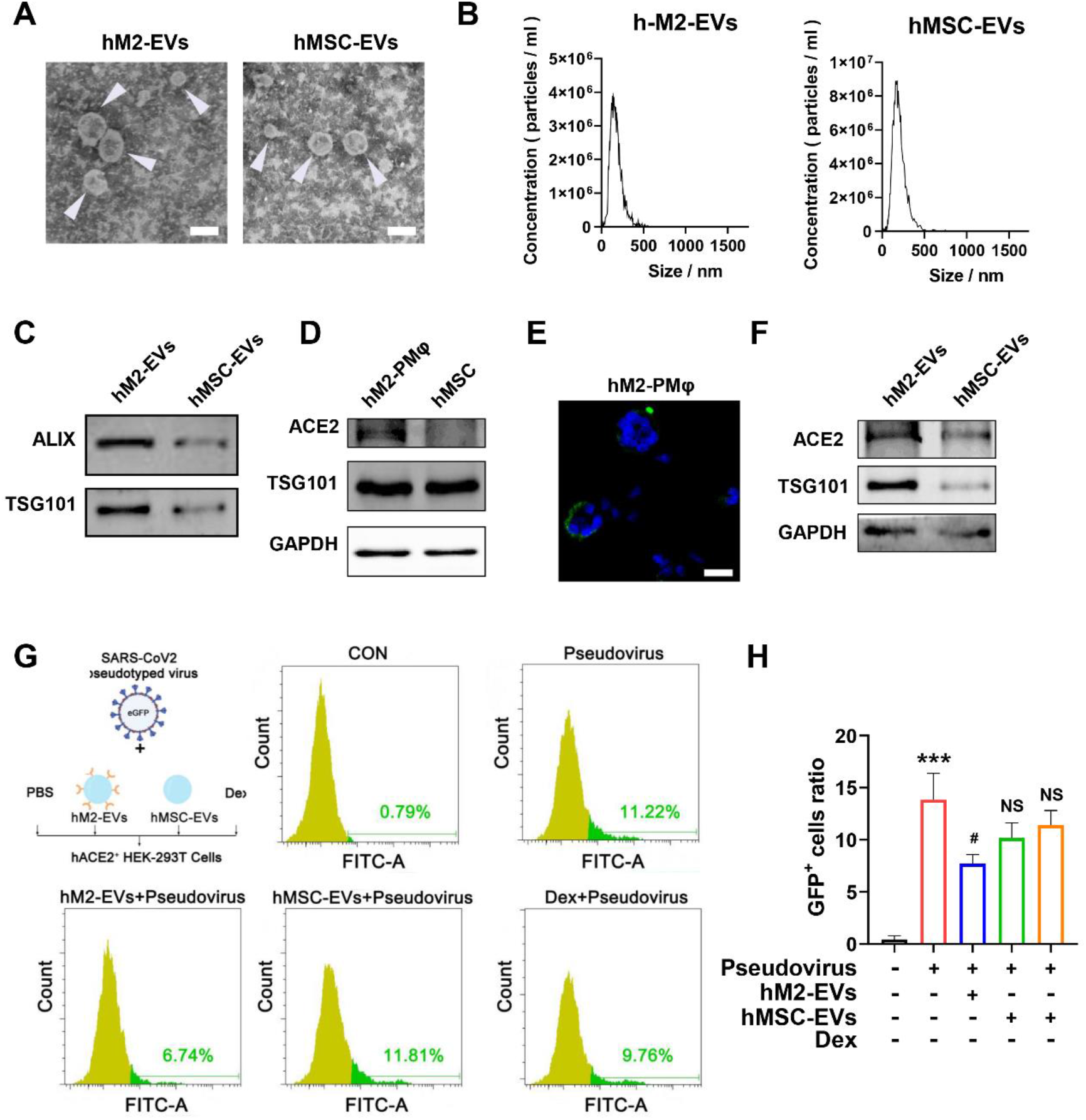
Effects of M2-EVs on SARS-CoV-2 pseudovirus infection *in vitro*. (A) Representative TEM images of human EVs. White arrow indicated EVs (scale bar = 200 nm). (B) Size distributions of human EVs measured by NTA. (C) Western blot analysis of positive marker proteins (ALIX and TSG101) in EV samples. (D) Western blot analysis of ACE2 protein expression in different cells. (E) Representative micrographs of ACE2 IF staining (scale bars = 20 μm). (F) Western blot analysis of ACE2 protein expression in different EVs. (G) FCA analysis of the percentage of infected GFP^+^ ACE2-overexpressed 293T cells after incubation with SARS-CoV-2 pseudovirus alone or in combination with hM2-EVs, hMSC-EVs or Dex for 48 h. (H) Quantification of the percentage of infected ACE2-overexpressed 293T cells in different groups (n = 3; ^***^p < 0. 001 *vs* CON group; ^#^p < 0.05 *vs* Pseudovirus group; ^NS^p > 0.05 *vs* Pseudovirus group).

Furthermore, we detected the expression of some important protein cargos (*e*.*g*., IL-10, TGF-β, HGF and SOD2) in M2-EVs, since these proteins have been associated with immunoregulatory or antioxidant effects of EVs[21, 35]. Indeed, we found the presence of these proteins within M2-EVs (Fig. 7F), suggesting that these functional proteins might also partly contribute to the therapeutic effects of M2-EVs. Altogether, these results suggest that M2-EVs can inhibit multiple proinflammatory pathways by regulating complicated miRNA-gene and gene-gene networks, and this effect was likely a joint result of many functional miRNAs and proteins delivered by these EVs.

### hM2-EVs served as nanodecoys to prevent SARS-CoV-2 pseudovirus infection *in vitro*

In addition to bacteria, other pathogens, such as many viruses, can also trigger cytokine storm. For example, in recent COVID-19 pandemic, the dramatic elevation of multiple cytokines, such as TNF-α, IL-6 and IFN-γ, were observed in the blood of patients with severe SARS-CoV-2 infection, and its severity was strongly associated with multiple organ injury and morality[37, 48]. Although some immunosuppressive drugs, such as Dex, had shown certain effects to reduce cytokine levels in COVID-19 patients[49]. However, the overall immunosuppression and possible side effects (such as hypertension and osteonecrosis of the femoral head) caused by these drugs, may adversely reduce the viral clearance and increase the risks of secondary infections [37, 48]. Thus, novel therapies that can prevent SARS-CoV-2 replication and its associated cytokine production are urgently needed. The key step of SARS-CoV-2 entrance of cells is primarily mediated by the binding of SARS-CoV-2 spike protein to the angiotensin-converting enzyme 2 (ACE2) receptor expressed on the cell surface[50]. Inspired by this process, the therapeutic strategies that aim to block or neutralize the binding of SARS-CoV-2 to ACE2 have been proposed. For example, EVs derived from ACE2-overexpressed parent cells have been used as nanodecoys to protect against pseudotyped SARS-CoV-2 infections [24, 51].

Interestingly, the expression of ACE2 receptor has been found on the surface of primary alveolar macrophages [52]. Based on these findings, we speculated that hM2-EVs may also express ACE2 and thus are capable of neutralizing SARS-CoV-2 virus. Our results showed that large amounts (∼25%) of human peritoneal macrophages (hPMφ) could be collected from peritoneal dialysate of PD patients (Fig. S7A). These cells were further polarized towards M2 phenotype *in vitro*, as indicated by elevated expression of Mrc1 and Arg1 (Fig. S7B-C). We isolated EVs from hPMφ and human MSCs, and the resulting hM2-EVs and hMSC-EVs were bi-layer lipid membrane vesicles with average sizes of ∼150 nm and ∼170 nm, respectively (Fig. 8A-B). Both EVs expressed markers such as Alix and TSG101 (Fig. 8C). We found that hM2-PMφ and hM2-EVs highly expressed ACE2, while a much weaker ACE2 expression was found in hMSC and hMSC-EVs, which was consistent with previous report that hMSCs had lower or negative ACE2 expression[53]. Next, we evaluated the potential of ACE2^+^ hM2-EVs in preventing SARS-CoV-2 pseudovirus infection *in vitro*. According to the previous report, ACE2-overexpresed 293T cells were established and then were incubated with SARS-CoV-2 pseudovirus containing a GFP reporter for detection of infected cells (Fig. S8A-C). Besides hM2-EVs, hMSC-EVs and Dex were also included in this experiment, since they had been proposed in the clinical trials of SARS-CoV-2 patients[54, 55]. As expected, elevated numbers of infected cells were observed in response to higher concentrations of SARS-CoV-2 pseudovirus (Fig. 8G-H). hM2-EVs treatment significantly reduced the numbers of infected cells compared to pseudovirus alone. However, neither hMSC-EVs nor Dex treatment could prevent pseudovirus infection in ACE2^+^ 293T cells (Fig. 8G-H). Although the *in vivo* viral infection experiment cannot be performed due to the absent of animal model and biosafety concerns, our results at least partly suggested that these ACE2^+^ hM2-EVs might be served as nanodecoys to neutralize SARS-CoV-2 and thus reduce viral load in ACE2^+^ cells. In addition, we found that ACE2^+^ hM2-EVs could also reduce cytokine release in mPMφ with LPS challenge (Fig. S7D), which was similar to mouse M2-EVs. Taken together, our results indicate that peritoneal M2-EVs can not only directly suppress cytokine production by delivering multiple functional cargos, but can also serve as nanodecoys to prevent SARS-CoV-2 infection because of their surface ACE2. Moreover, as a type of cell-derived nanomaterial, the therapeutic potential of M2-EVs can be readily and precisely improved by genetic and chemical modifications, and loading with specific therapeutic reagents in response to different diseased conditions. Therefore, M2-EVs may be a promising multi-target nanomedicine to attenuate severe infections-related cytokine storm, such as recent COVID-19.

## Conclusion

In summary, primary peritoneal M2 macrophages exhibited superior anti-inflammatory potential than immobilized cell lines in decreasing bacterial endotoxin-induced cytokine (e.g., TNF-α, IL-6) release. M2-EVs were able to decrease bacterial endotoxin-induced cytokine (*e*.*g*., TNF-α, IL-6) release and its associated oxidative stress and multiple organ damage, which might be mediated by inhibiting multiple key proinflammatory pathways (*e*.*g*., NF-κB, JAK-STAT, p38 MAPK) *via* delivering a set of functional cargos and regulating complex miRNA-gene/gene-gene networks, which is different from the existing single cytokine-targeted therapy. Besides, human peritoneal M2-EVs were able to exert as anti-inflammatory agents and serve as nanodecoys to prevent SARS-CoV-2 pseudovirus infection *in vitro* due to their surface angiotensin-converting enzyme 2 (ACE2), a receptor of SARS-CoV-2 spike protein.

## Supporting information

Manuscript

## Acknowledgments

This work was partly supported by the National Natural Science Foundation of China (32071453, 31871001, 81571808, 81774160), Health Commission of Sichuan Province (20PJ003), and 1.3.5 Project for Disciplines of Excellence (ZYGD18014), West China Hospital of Sichuan University. The authors would like to thank Sichuan Neo-life Stem Cell Biotech &Sichuan Stem Cell Bank for providing the hMSCs, Jie Zhang and Lin Bai for providing technical assistance with confocal microscopy, and Shisheng Wang for providing technical assistance with the bioinformatics analysis.

## Author contributions

Jingping Liu and Yizhuo Wang designed the project. Jingping Liu, Yizhuo Wang, Shuyun Liu, Peng Lou, Meng Zhao, Lan Li, Ke Lv, Yujia Yuan and Meihua Wan performed the experiments, collected the data, and analyzed and interpreted the data. Ling Li and Xueli Zhou collected the patient peritoneal dialysate samples. All authors contributed to the writing of the manuscript, discussed the results and implications, and edited the manuscript at all stages.

