## Supplementary material for "Peritoneal M2 macrophage-derived extracellular vesicles as natural multi-target nanotherapeutics to attenuate cytokine storm after severe infections": Manuscript

### Supplementary methods

#### Collecltion and concentration of culture medium

The culture medium of M2-PMφ or M2-RAW264.7 cells was collected and then concentrated using ultrafiltration tubes (3 Kda cut-off, Millipore). The protein amounts of concentrated medium were determined by BCA kit and normalized to the cells numbers in each group. The concentrated culture medium was stored at -20℃ for further use.

#### Detection of intracellular mitochondiral ROS (mtROS)

The levels of intracellular mtROS were measured by flow cytometry analysis. After treatment with indicated conditions, cells were incubated with MitoSOX (2 μM, Thermo Fisher Scientific, Sunnyvale, CA, USA) for 30 min, washed twice with PBS and analyzed using a flow cytometer (Beckman, USA).

**Supplementary Figures and Tables**

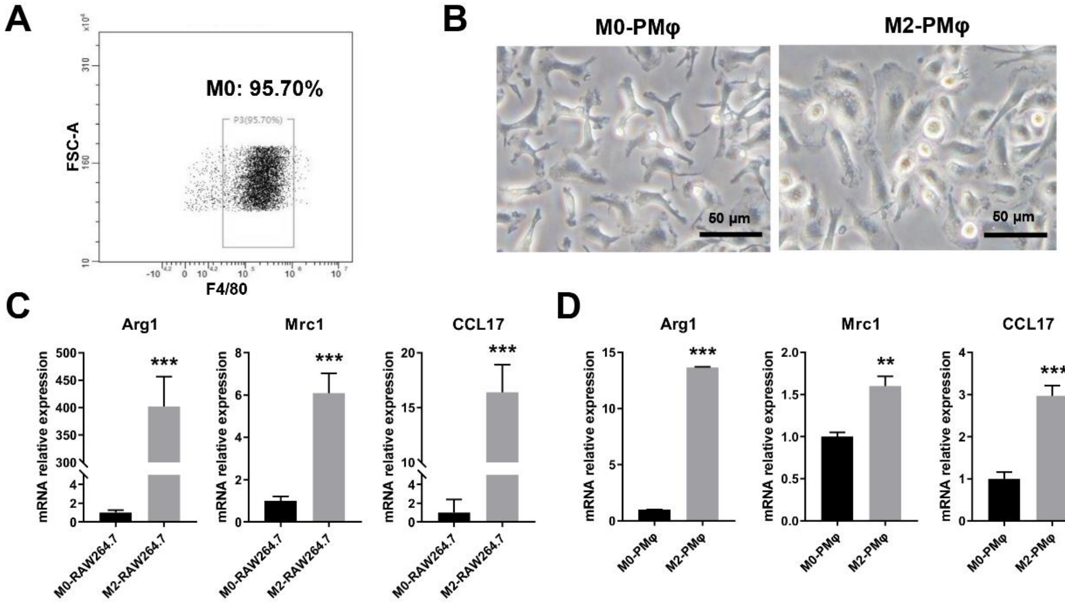


**Fig. S1 Polarization and characterization of mouse M2 macrophages.** (A) FCA analysis of the purity isolated PMφ. (B) Representative light field images of M0 and M2 PMφ (scale bar = 50 μm). (C) Measurement of Arg1, Mrc1 and CCL17 gene expressions in IL-4/IL-13-induced M2-RAW264.7 cells for 48 h by qPCR. (D) Measurement of Arg1, Mrc1 and CCL17 gene expressions in IL-4/IL-13-induced M2-PMφ cells for 48 h by qPCR. (n = 3; $^{***}p < 0.001$ *vs.* M0 group)

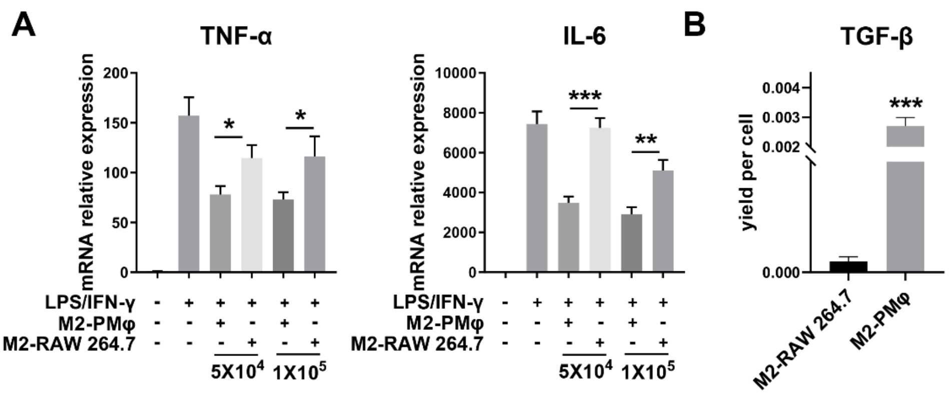


**Fig. S2 Effects of M2-PMφ and M2-RAW264.7 on inflammatory response *in vitro*.** (A) Measurement of TNF-α and IL-6 mRNA levels in LPS/IFN-γ primed PMφ with or without the concentrated culture for 4 h by qPCR. (B) Measurement of TGF-β levels of cultures from M2-Mφ by ELISA. (n = 3; $^{***}p < 0.01$, $^{**}p < 0.01$, $^{*}p < 0.05$ M2-PMφ group *vs* M2-RAW264.7 group)

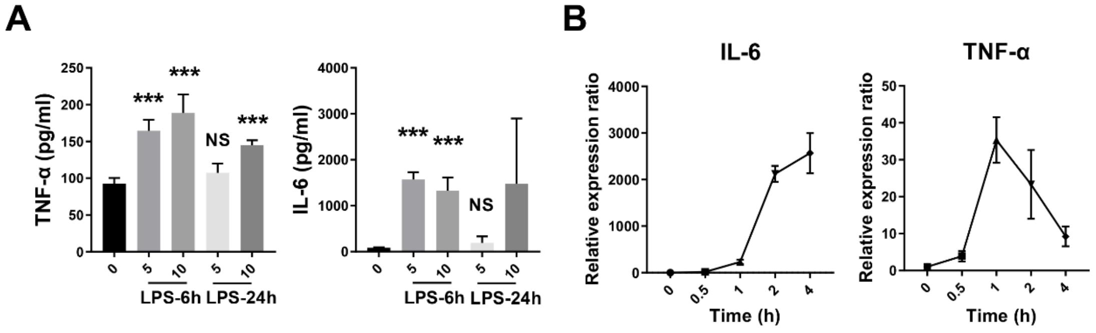


**Fig. S3 Establishment and evaluation of LPS-induced cytokine storm mouse model.** (A) Measurement of mouse plasma TNF-α and IL-6 levels at 6 h and 24 h after LPS (5, 10 mg/kg) challenge by ELISA (n = 3; $^{***}p < 0.001$, $^{NS}p > 0.05$ *vs.* CON group). (B) The dynamic changes in the plasma TNF-α and IL-6 levels at 0.5 h, 1 h, 2 h and 4 h after LPS challenge (n = 5).

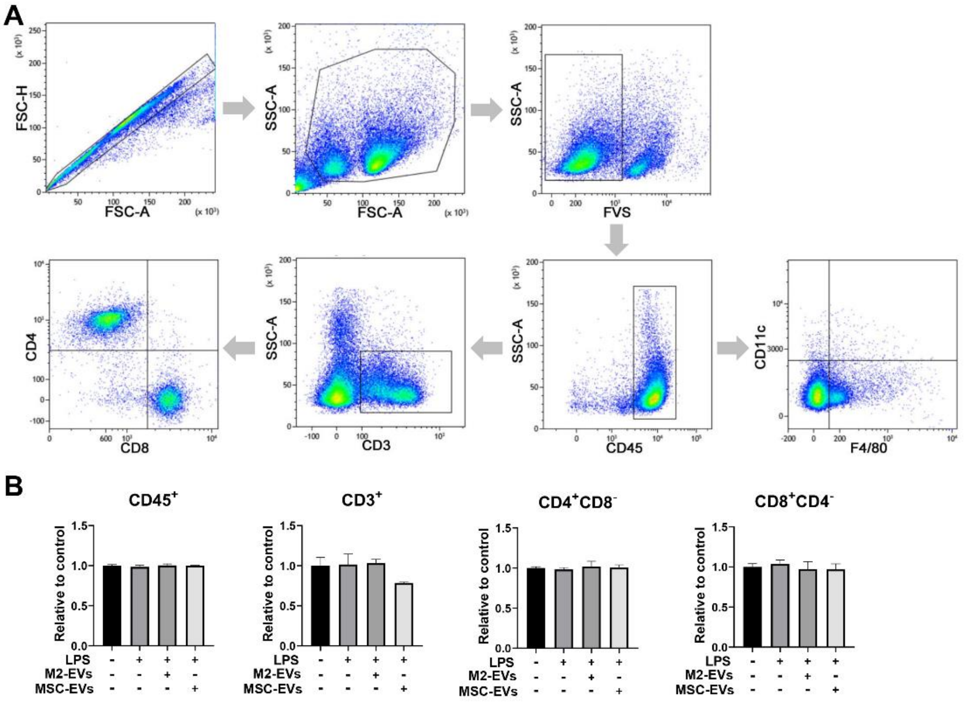


**Fig. S4 FCA analysis of immune cells in tissue samples of mice.** (A) FCA gating strategies for the sorting of different types of immune cells in spleen tissues. (B) Quantification the populations of $CD45^{+}$ total leucocytes, $CD3^{+}$ total T cells, $CD4^{+}$ helper T cells or $CD8^{+}$ cytotoxic T cells population in spleen tissues of each group.

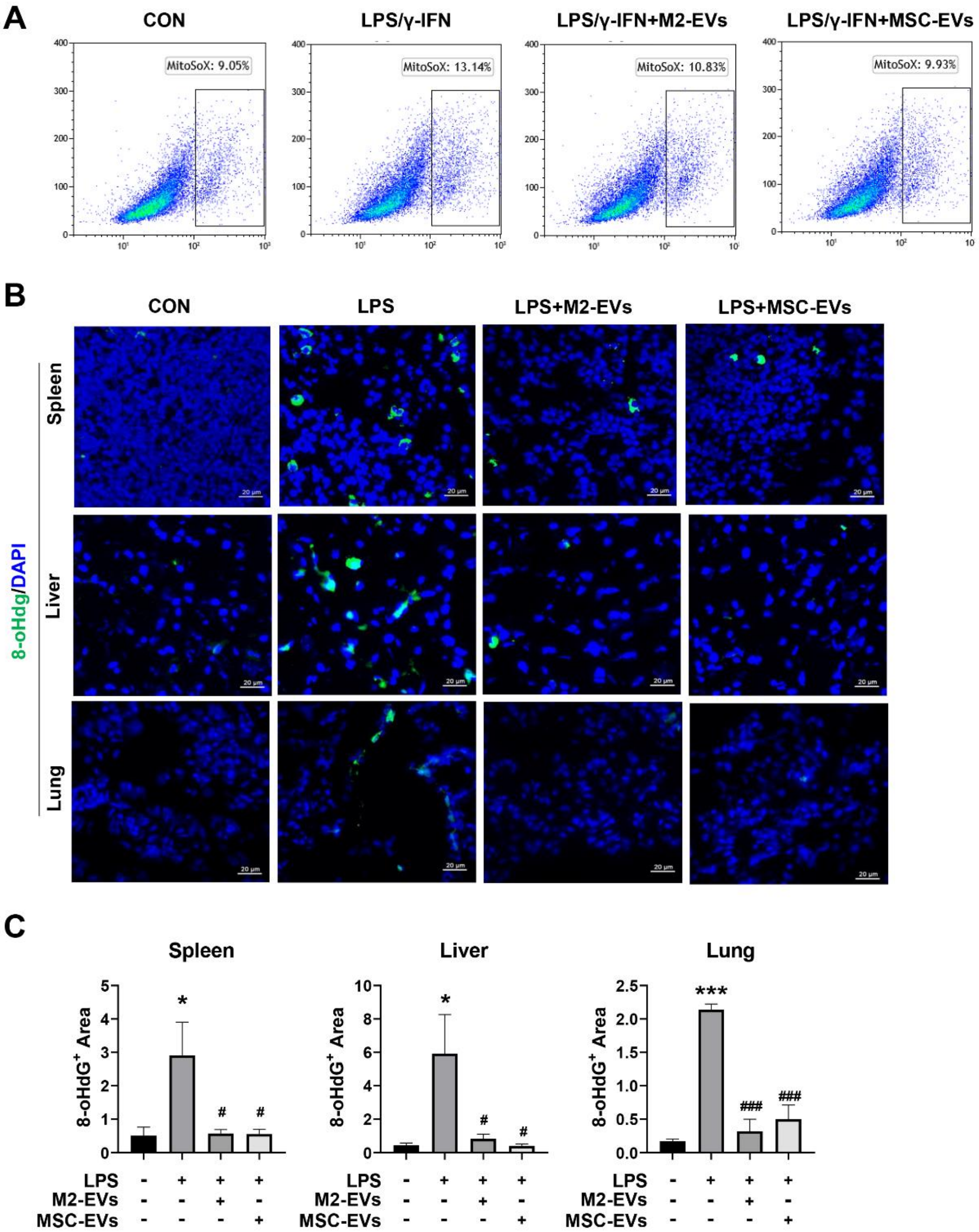


**Fig. S5 Effects of M2-EVs treatment on oxidative stress *in vitro* and *in vivo*.** (A) FCA analysis of mtROS levels of LPS/IFN-γ-primed PMφ with or without different EVs treatments for 4 h using MitoSOX staining. (B) Representative micrographs of 8-OHdG IF staining of spleen, liver and lung sections at 24 h after LPS challenge with or without EVs treatments (scale bars = 20 μm). (C) Quantification of 8-OHdG$^{+}$ cells in the lung, liver and spleen sections detected by IF staining (n = 3; $^{***}p < 0.001$, $^{*}p < 0.05$ *vs.* CON group; $^{\#\#\#}p < 0.001$, $^{\#}p < 0.05$ *vs.* LPS group).

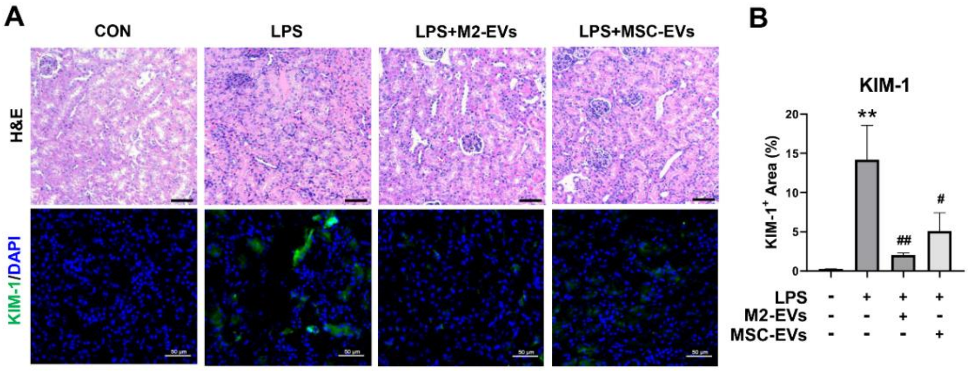


**Fig. S6 Effects of M2-EVs treatment on renal injury *in vivo*.** (A) Representative micrographs of H&E staining and KIM-1 IF staining of the kidney sections at 24 h after LPS challenge with or without different EVs treatments (scale bar = 50 μm). (B) Quantification of KIM-$1^+$ cells in the kidney detected by IF staining (n = 3; $^{**}p < 0.01$ *vs*. CON group; $^{\#}p < 0.05$, $^{\#\#}p < 0.01$ *vs*. LPS group).

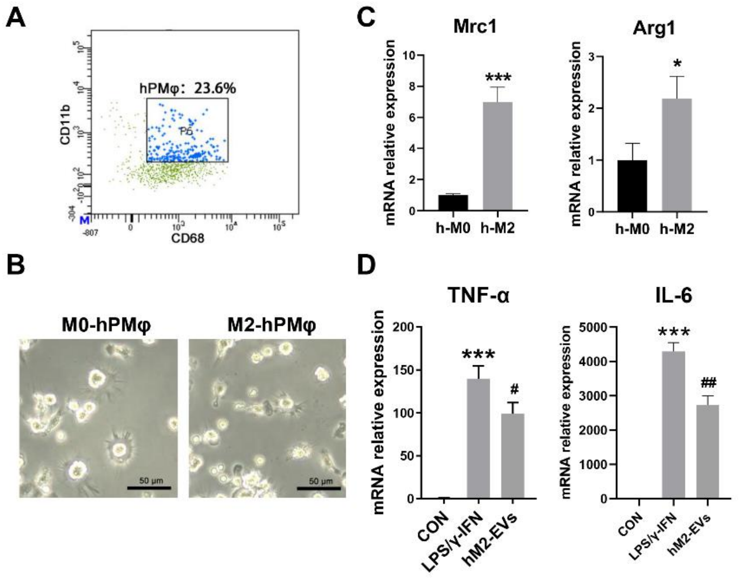


**Fig. S7 Isolation and polarization of human peritoneal M2 macrophages *in vitro*.** (A) Representative light field images of M0-hPMφ and M2-hPMφ (scale bar = 50 μm). (B) FCA analysis of the purity of isolated hPMφ. (C) Quantification of Mrc1 and Arg1 mRNA levels in IL-4/IL-13-induced M2 hPMφ for 48 h by qPCR. (n = 3; $^{***}p < 0.001$, $^{*}p < 0.05$ *vs* M0-hPMφ). (D) Measurement of TNF-α and IL-6 mRNA levels in LPS/IFN-γ primed PMφ with or without hM2-EVs treatment for 4 h by qPCR. (n = 3; $^{*}p < 0.05$).

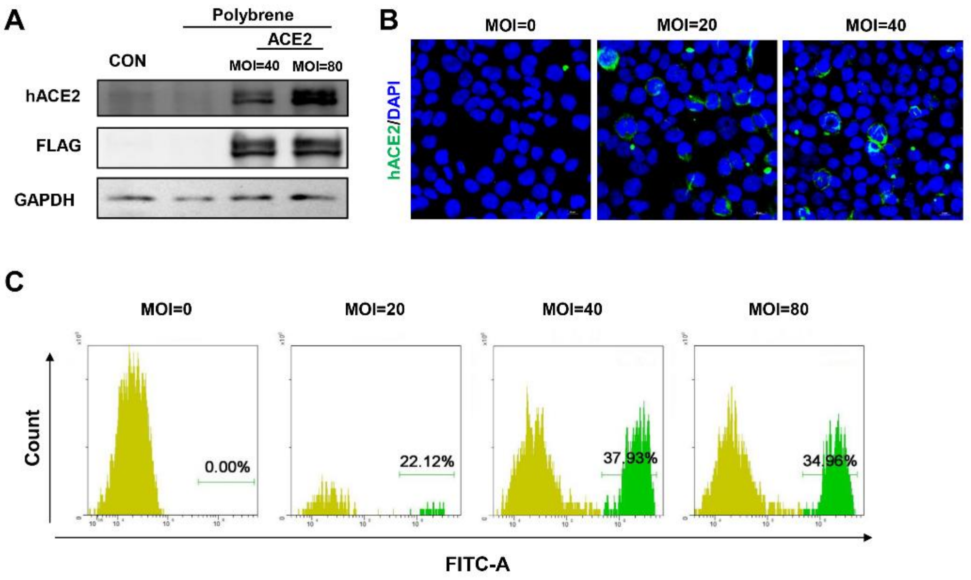


**Fig. S8 Establishment of SARS-CoV-2 pseudovirus infection model *in vitro*.** (A) Western blot analysis of ACE2 expression in 293T cells transfected with ACE2 lentivirus for 48 h with different MOI. (B) Representative micrographs of ACE2 IF staining in 293T cells transfected with ACE2 lentivirus for 48 h with different MOI. (C) FCA analysis of the percentage of infected $GFP^+$ ACE2-overexpressed 293T cells after incubation with different MOI of SARS-CoV-2 pseudovirus.

**Table S1. The primer sequences used in this study.**

| | |
|---|---|
| mTNF-α- Forward | ACGGCATGGATCTCAAAGAC |
| mTNF-α-Reverse | AGATAGCAAATCGGCTGACG |
| mIL-6-Forward | GTTCTCTGGGAAATCGTGGA |
| mIL-6-Reverse | TGTACTCCAGGTAGCTATGG |
| mArg1-Forward | AGACAGCAGAGGAGGTGAAGAG |
| mArg1-Reverse | CGAAGCAAGCCAAGGTTAAAGC |
| mMrc1-Forward | GTCTGAGTGTACGCAGTGGTTGG |
| mMrc1-Reverse | TCTGATGATGGACTTCCTGGTAGCC |
| mCCL17-Forward | GAGCCATTCCCCTTAGAAAG |
| mCCL17-Reverse | AGGCTTCAAGACCTCTCAAG |
| mTGF-β1-Forward | CAACAATTCCTGGCGTTACCTTGG |
| mTGF-β1-Reverse | GAAAGCCCTGTATTCCGTCTCCTT |
| hMrc1-Forward | GACGTGGCTGTGGATAAATAAC |
| hMrc1-Reverse | CAGAAGACGCATGTAAAGCTAC |
| hArg1-Forward | GACCTGCCCTTTGCTGACATCC |
| hArg1-Reverse | TCTTCTTGACTTCTGCCACCTTGC |
